# Y recombination arrest and degeneration in the absence of sexual dimorphism

**DOI:** 10.1101/2021.05.18.444606

**Authors:** Thomas Lenormand, Denis Roze

## Abstract

Current theory proposes degenerated sex chromosomes evolve via three successive steps: recombination arrest, which links male-beneficial alleles to the Y chromosome; degeneration of these regions due to the inefficacy of natural selection in the absence of recombination; and lastly, the evolution of dosage compensation to correct the resulting low expression of X-linked genes in males. Here we investigate new models of sex chromosome evolution incorporating the coevolution of cis- and trans-regulators of gene expression. We show that the early emergence of dosage compensation favors the maintenance of Y-linked inversions by creating sex-antagonistic regulatory effects. This is followed by inversion degeneration caused by regulatory divergence between the X and Y chromosomes. In stark contrast to the current theory, the whole process occurs without any selective pressure related to sexual dimorphism.

**One Sentence Summary:** Turning sex chromosome theory on its head: early evolution of dosage compensation can maintain successively forming Y chromosome strata that undergo genetic degeneration.

## Main Text

Many species have XX/XY chromosomal sex determination systems (*1*). Y chromosomes are often non-recombining and degenerated. Suppressed genetic recombination initiates genetic degeneration, and Y chromosome evolution of an autosomal ancestor. In several chiasmate species, recombination suppression has been shown to involve successive events, each affecting Y sub-regions of different sizes, called “strata”, that are detected from differences in sequence divergence from the homologous X regions (*2*–*5*). Current theory proposes that Y chromosomes evolve through three steps. Step 1 is a consequence of the evolution of sexual dimorphism (*9*): divergent selection in males and females may generate intralocus sexual conflict, which, for loci linked to a sex-determining locus, favours suppressed recombination, so that the allele favorable in one sex becomes associated with that sex (*6*–*13*). In the second step, selective interference in the absence of recombination reduces the efficacy of natural selection, leading to accumulation of deleterious mutations on the Y and genetic degeneration. Finally, the model proposes a third step in which dosage compensation evolves to restore optimal gene expression in males, whose sex-linked genes have lowered expression due to degeneration (*7, 14*–*16*). The compensation process involve various mechanisms in different species, and compensation is not always complete for all X linked genes (*17*–*20*). The steps in this theory have been studied over the past c.a. 50 years, both empirically and theoretically (*5, 6,21*–*25*). Empirical support for the first step is particularly equivocal where despite decades of investigation (*3, 5, 26, 27*), decisive evidence for a causal role of sexually-antagonistic loci on recombination arrest, is conspicuously lacking. The second step is difficult to reconcile with the observation of small degenerated strata (*5*), within which selective interference should be minimal. Lastly, the causal ordering of events has also been challenged by observation of the early evolution of partial dosage compensation in young sex-chromosomes (*28*– *32*). Theoretically, each step suffers from well-identified limitations (sup. mat. 1). However, an important global limitation is that each step has generally been considered independently from the others, resulting in a piecemeal set of models lacking integration. In particular, regulatory changes have not been consistently studied throughout the whole process. Yet, they can influence the evolution of sex-limited expression in the first step, they can contribute to compensatory adaptive silencing in the second step, and they are pivotal for the evolution of dosage compensation in the third step.

This paper investigates a new model that includes the joint evolution of regulatory changes and accumulation of deleterious mutations, and can lead to the evolution of an autosome into a degenerated sex chromosome with dosage-compensation. We use individual-based stochastic simulations of a population of *N*_*pop*_ diploid individuals, with XY males and XX females (sup. mat., Fig S6). We start by considering the evolution of a pair of autosomes carrying hundreds of genes subject to partially recessive deleterious mutations, with one homolog having newly acquired a sex-determining locus. Gene expression is controlled by cis-regulatory sequences (affecting expression on the same chromosome as themselves) interacting with trans-regulators affecting the gene copies on both homologs (*33*). All these elements can mutate. To allow for possible dosage compensation, we assume that each gene is controlled by one male-and one female-expressed trans-regulator (Fig S6). We assume that each gene’s overall expression level is under stabilizing selection around an optimal level and that the relative expression of the two copies of each gene determines the dominance level of the deleterious mutations occurring in the gene. For instance, a deleterious mutation occurring in a relatively less expressed gene copy will be less harmful than one in a relatively more expressed copy. We assume that mutations occur that suppress recombination on a segment of the Y. We refer to these mutations as inversions for simplicity, although they could correspond to other mechanisms of recombination arrest (sup. mat. 2). Inversions of any size can occur, but we follow only those on the Y that include the sex-determining locus, which will necessarily be confined to males and cause recombination arrest. We assume that inversions can “add-up”, meaning that new inversions can occur on chromosomes already carrying a previous inversion, and thus extend the non-recombining part of the Y. Finally, we assume that reversions restoring recombination can occur, and, for simplicity, that such reversions cancel only the most recent inversion (see sup. mat. 2 for justifications of these assumptions).

To understand the dynamics of sex chromosome evolution in this model, it is useful to first consider the case where the cis- and trans-regulators do not mutate. In this case, all inversions on the Y are eventually reversed and lost. This occurs in two steps. First, an inversion appears on a given Y and “freezes” a segment of the chromosome. If, by chance, this Y carries relatively fewer or milder deleterious mutations, this “lucky” inversion has a selective advantage and consequently tends to fix in the population. This process has been described for autosomes by Nei (*34*), but an important difference here is that a fixed Y-linked inversion stays heterozygous in males, and therefore causes recombination arrest. In contrast, a fixed autosomal inversion is homozygous and does not cause recombination arrest. Larger inversions are overrepresented among these lucky inversions, as they contain more genes and exhibit a larger fitness variance (Fig 1A, sup. mat.1). However, after fixation among the male-determining chromosomes in the population, they start accumulating deleterious mutations due to selective interference. Fitness declines faster for larger inversions due to stronger selective interference (Fig 1C). When the inversion’s marginal fitness becomes lower than the fitness of the corresponding chromosomal segment on the X, reversions are selectively favored and spread, which restores recombination. Thus, Y-specific inversions are maintained only transiently in the population in the absence of regulatory mutations (Fig 1B). These periods of recombination suppression do not last long enough to lead to Y degeneration.

**Fig 1.**
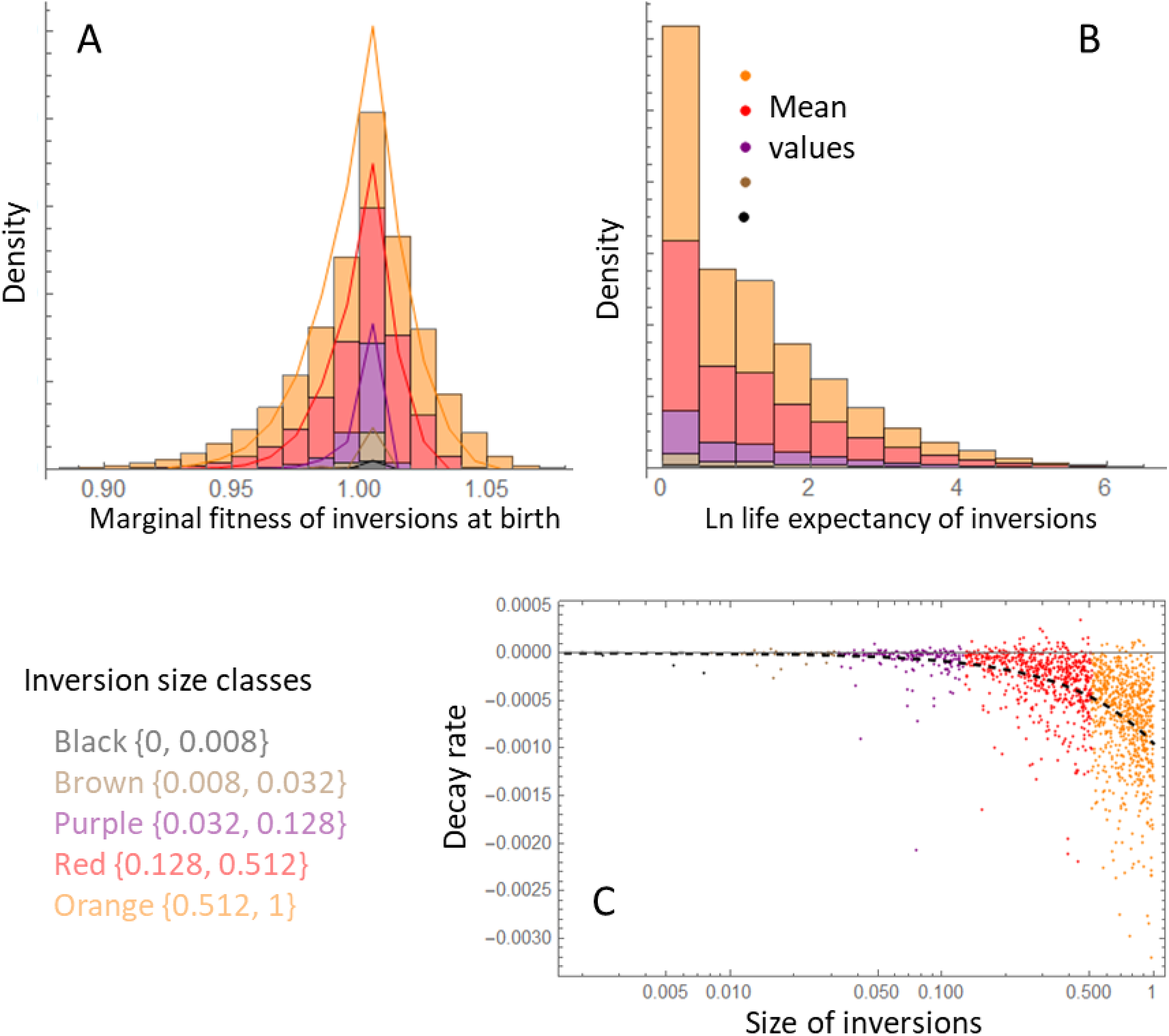
Characteristics of inversions in the absence of regulatory evolution. **(A)** Cumulated distributions of the marginal fitness of newly arising inversions for different size classes of inversions (the different colors, legend at bottom left; the whole chromosome has size 1 on this scale, the sex-determining locus being located at one end of the chromosome). When an inversion arises, the corresponding chromosomal segment carries by chance more or less deleterious mutations than average, resulting in a variance in marginal fitness. Lines: expectations computed assuming a Poisson distribution of the number of mutations per inversion, each with a fixed effect *s* = *s*_*mean*_). **(B)** Distribution of the log time before inversions become extinct for the different size classes (in number of generations). These distributions have approximately the same mean values (indicated by the corresponding color dots, on an arbitrary y-scale). **(C)** Marginal fitness decay rate per generation for inversions (y-axis), as a function of their size (x-axis, log scale). Inversions accumulate deleterious mutations because of selective interference. This decay rate is computed over the first 50 generations on the relative marginal fitness of newly arising inversions. Only inversions lasting at least 50 generations are therefore represented. The dashed line is the least square fit of a power law yielding *y* = −0.00095*x*^1.06^, indicating that this decay rate varies approximately linearly with inversion size. Parameter values are described in methods (*N*_*pop*_ = 10^4^, *I* = 0.1), but with no mutation on regulators (*U*_*c*_ = *U*_*t*_ = 0).

A radically different four-step process emerges when the regulatory sequences can mutate and evolve. A typical example is illustrated in Fig 2. It starts, as before, with the fixation of a lucky inversion on the Y. However, once the inversion stops recombination, X and Y cis-regulators can start evolving independently (step 2). We showed previously that this creates a positive feedback loop that causes rapid degeneration of Y-linked alleles (*35*): by chance, some genes on the Y become slightly less expressed than their X-linked alleles, and accumulate more deleterious mutations (because lower expression makes mutations more recessive), selecting for a further reduction of Y expression to make them even more recessive. This process can work on individual genes, irrespective of the size of the non-recombining region created by the inversion (*35*), and the degeneration does not involve selective interference. However, like in the absence of regulator evolution, recombination arrest also triggers the accumulation of deleterious mutations by selective interference, especially if the inversion includes many genes.

**Fig 2.**
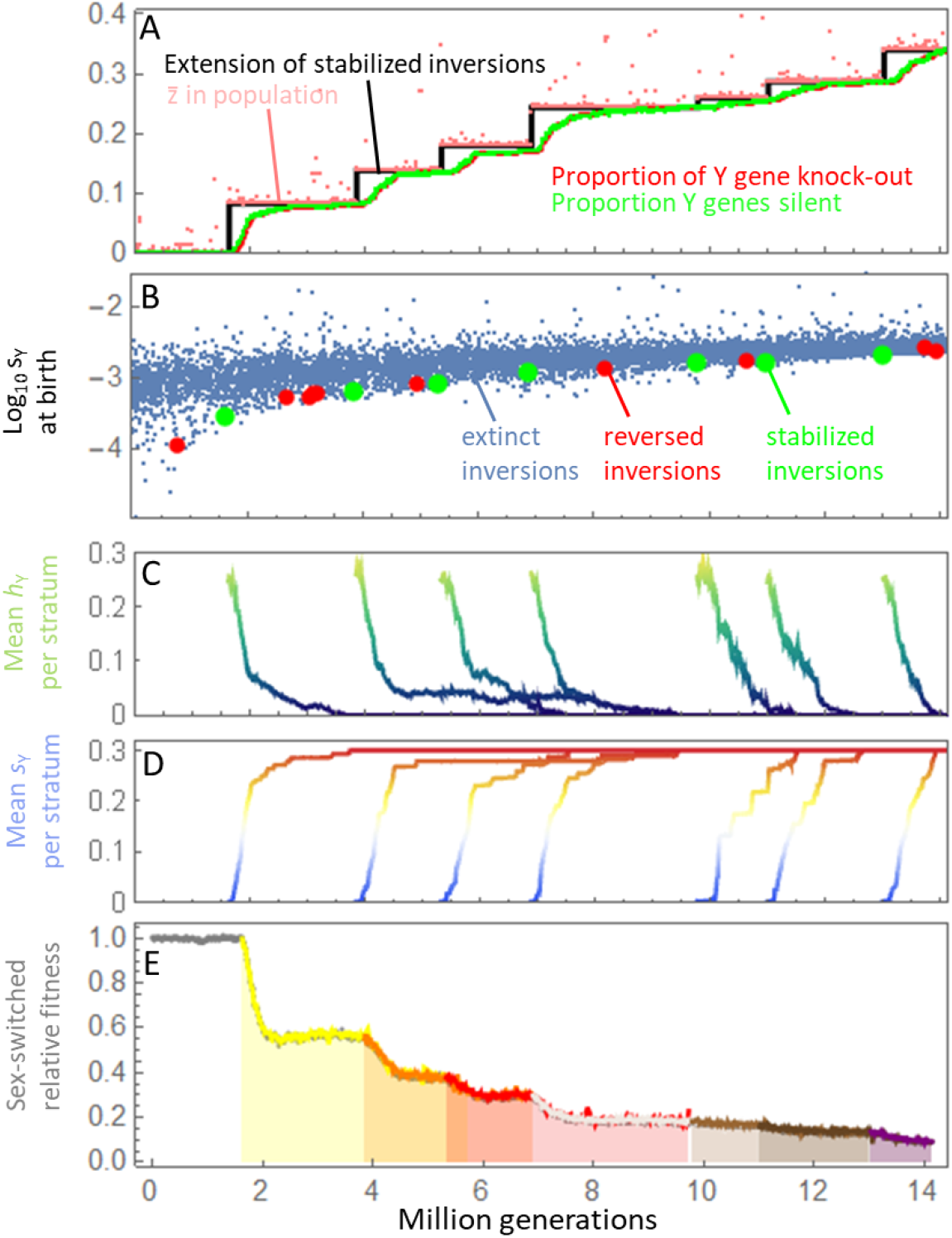
Example of a typical Y degeneration process. The Y progressively degenerates by the accumulation of inversions, which accumulate deleterious mutations, evolve dosage compensation with sex-antagonistic fitness effects, and become immune to reversions. **(A)** Black stairplot: extension of each successive stratum of the Y (expressed as the fraction of the physical length of the Y), corresponding to stabilized inversions. Pink dots : average fraction of the non-recombining Y in the population 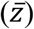. Green: proportion of Y genes that are silenced. Red: proportion of Y genes that accumulate deleterious mutations effects up to *s*_*max*_ (here *s*_*max*_ = 0.3), corresponding to a gene knock out. **(B)** Log_10_ of the average effect of deleterious mutations carried by inversions when they first arise in the population (averaged over the different genes within the inversion). Blue dots: random subsample of inversions that are eventually lost before fixing in the population. Red dots: inversions that reach fixation, but become extinct after the occurrence of a reversion. Green dots: inversions that reach fixation, and last until the end of the simulation (stabilized inversions), becoming strata on the Y. **(C)** Mean dominance of deleterious mutations on each stabilized inversion (initial dominance of deleterious mutations is 0.25). **(D)** Accumulation of deleterious mutation on each stabilized inversion (the maximum effect *s*_*max*_, is set to 0.3 for all genes). **(E)** Fitness that the Y carrying the stabilized inversions would have on average, if expressed in a female (relative to the actual average fitness of males). The different colors highlight the occurrence of the successive strata. The average fitness of males that would carry two X chromosomes at that time is indicated in gray, but yields the same values. Parameter values are described in the methods section (*N*_*pop*_ = 10^4^, *I* = 0.1).

The key step is the third, which stabilizes inversions in the long term, even when they become entirely degenerated (Fig 3, Fig S4). Cis-regulator divergence and degeneration in step 2 cause a departure from optimal expression levels in males. Assuming that gene expression is under stabilizing selection, this causes sex-specific trans-regulators to diverge to correct this and maintain optimal expression in both sexes. For instance, if a Y cis-regulator weakens, causing lower expression, this will favour a stronger allele of the male trans-regulator, to compensate for the low expression by increasing expression (from both alleles of the gene). The divergence of X- and Y-linked *cis*-regulators, and the divergence of sex-limited trans-regulators, automatically generate sexually-antagonistic fitness effects: X *cis*-regulators that recombine onto the Y would cause overexpression in males (due to mismatches with male trans-regulators), and similarly Y *cis*-regulators recombined onto the X would cause under-expression in females. Hence, if a reversion occurs, the reestablished recombination between X and Y would reduce offspring fitness by creating a mismatch between *cis* and *trans* regulators. This sexually antagonistic effect caused by nascent dosage compensation protects diverging inversions from reversion. This is the ultimate cause of Y recombination suppression in our model (sup. mat. 3).

**Fig 3.**
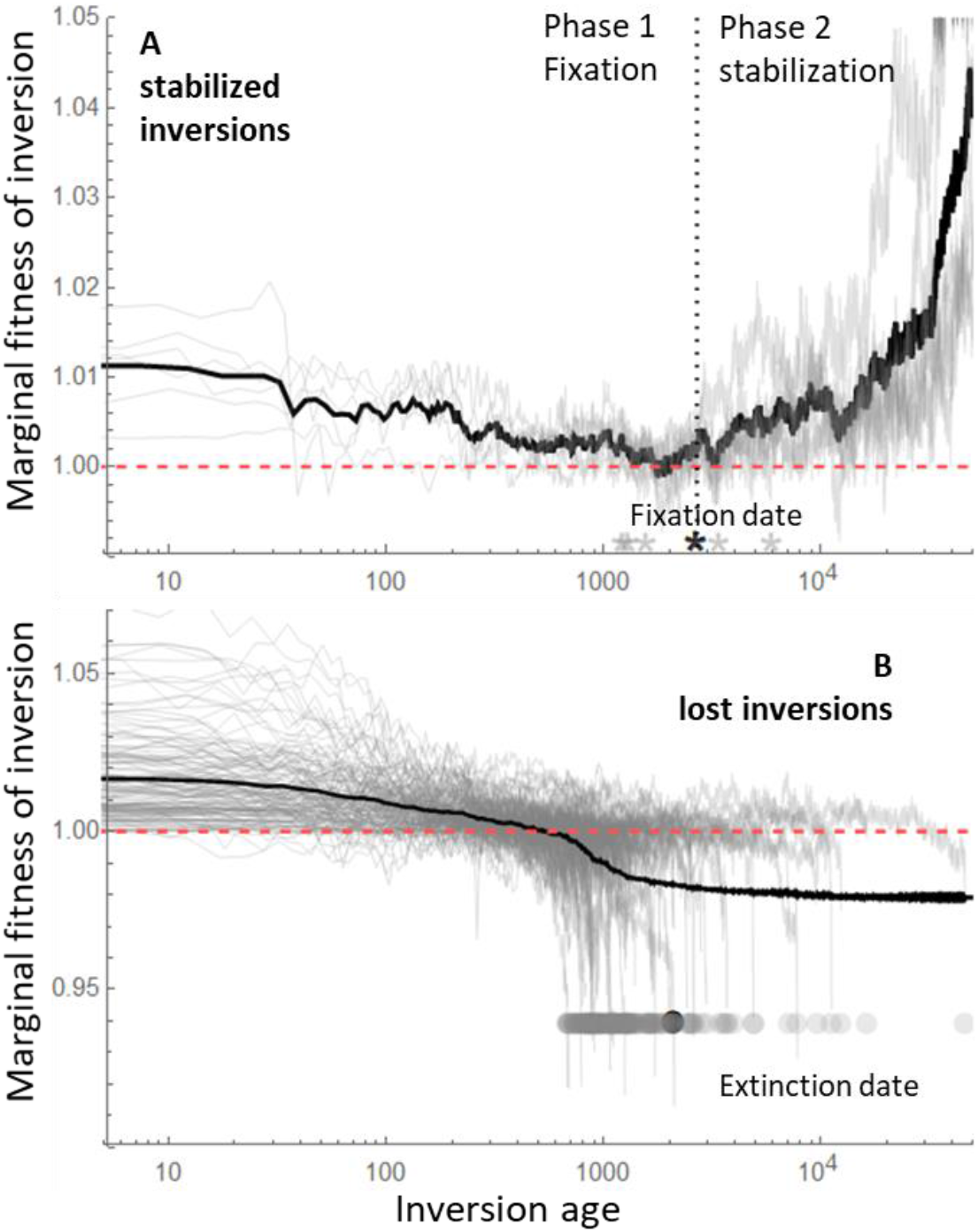
Fitness trajectories of stabilized and lost inversions. *x*-axis, inversion age: the number of generations since the appearance of the inversion (in log-scale). *y*-axis : marginal fitness of the inversion relative to the same chromosomal segment on the X if it was in a male (*W*_*margX*_, see methods). After fixation, this measures the sexually-antagonistic effect of nascent dosage compensation. The marginal fitness of the inversion relative to the same chromosomal segment among Y chromosomes not carrying the inversion (*W*_*margY*_, see method) yields undistinguishable results before the inversion fixes and is therefore not shown to not overload the figure (note that *W*_*margY*_ cannot be computed after the inversion fixes as all Y chromosomes carry the inversion). In gray, individual trajectories, in black average values. **(A)** Inversions that are stabilized as first Y strata, collected over 10 evolutionary replicates after 1 million generation. Their fixation date is indicated by a star at the bottom. (**B**) Inversions that are in the top 15 longest lived ones before a first stratum is stabilized, collected over 10 evolutionary replicates over 1 million generation. Their extinction date is indicated by a gray disk at the bottom. The time averaged fitness at time *t* (in black) is computed over all inversions, counting their last achieved fitness if they are extinct at *t*. Red dashed line indicates value 1.

Of course, only a minority of inversions evolve this nascent dosage compensation fast enough (relative to the speed of degeneration) to stay immune to reversion. However, a positive feedback loop is also operating here: when an inversion starts evolving dosage compensation, it becomes relatively immune to reversion and can maintain itself longer in the population, giving it more time to evolve further dosage compensation. The inversion becomes completely degenerated in the final fourth step, and has complete dosage compensation (at least for dosage-sensitive genes). This leads to very strong sexually-antagonistic regulatory effects, which effectively make the inversion immune to reversions. Degeneration may also lead to loss of homology between the Y and X, definitively removing the possibility of X-Y recombination.

In this model, recombination suppression evolves along with regulatory evolution, and, paradoxically, selective interference opposes it and therefore tends to slow down the process. The evolution of nascent dosage compensation involves the fixation of compensatory mutations and is partly adaptive. If interference is too strong, inversions accumulate deleterious mutations too fast and are quickly replaced by reversions. Accordingly, stabilized inversions tend to be heavily biased towards small sizes, though less so when population size is larger (Fig S1C). In large populations, recombination suppression and degeneration evolve more quickly, since more inversions occur and selective interference is relatively less efficient at removing large inversions (Fig S1). Finally, as expected, this overall process is faster when the intensity of stabilizing selection on gene expression levels is strong, since this effect on fitness is the key to the evolution of dosage compensation, selectively protecting partially degenerated inversions from reversions (Fig S2).

This theory suggests that the Y chromosome is entangled in an inescapable regulatory trap leading to recombination arrest and degeneration even in the absence of any selective pressure related to sex-dimorphism. Indeed, unlike the current theory (*10, 11, 36, 37*), our model only includes genes with the same optimal expression level in males and females, and deleterious mutations that have the same effect in both sexes. This process is inherently stochastic, as it involves the rare stabilization of a handful of inversions (Fig S3 shows the high variance of the process). However, it works faster in larger populations, as selective interference (whose effect is stronger in smaller populations) opposes recombination arrest and the stabilization of large strata. It also turns current theory on its head by showing that dosage compensation can cause recombination suppression, rather than being a consequence of degeneration after such suppression. Sexually-antagonistic effects are involved in the evolution of suppressed recombination. However, they result from the fact that one sex is heterogametic, not from males and females having divergent sex-specific optima for reproductive traits or expression levels. All genes whose dosage affects fitness can contribute to the process, not just a subset of sexually-antagonistic loci. The potential sexually-antagonistic effect of dosage compensation has long been appreciated (*7, 10, 11,15, 16, 23, 38, 39*). However, its potential role in recombination arrest has not been previously recognized, as it is usually thought to occur very late in the degeneration process. Once recombination has stopped, sexually-antagonistic alleles can easily occur and be maintained (*10, 11, 40*), but as shown here, they are not required for recombination arrest.

Overall, we showed that the emergence of a non-recombining and degenerated sex chromosomes in diploid organisms requires very few ingredients: genetic sex determination, deleterious mutations, inversions, sex-specific trans-regulators, and stabilizing selection on gene expression levels (Fig S5). This theory includes all steps in a single set of assumptions and is compatible with current data on sex chromosome evolution in chiasmate species (sup. mat. 3): notably the occurrence of strata, including small ones (reviewed by *5*), the occurrence of early regulatory changes in young sex-chromosomes (*28*–*32*) and the lack of decisive evidence for a causal role of sexually-antagonistic loci on recombination arrest, despite decades of investigation (*3,5,26, 27*).

## Acknowledgments

We thank D. Charlesworth, G. Marais, Y. Michalakis for comments and suggestions, and K McKean for editing. We thank MBB cluster from Labex CEMEB, and CNRS ABiMs cluster.

## Funding

This work was supported by grant GenAsex ANR-17-CE02-0016-01.

## Author contributions

Original idea TL, DR; Model conception TL, DR; Code DR, TL; Simulations TL; Data analyses TL; Interpretation TL, DR; First draft, editing and revisions TL, DR; Project management and funding TL.

## Competing interests

the authors declare no conflict of interest.

## Data and materials availability

simulation code will be available after publication on GitHub.

## Methods

### 1. Model

#### Genome

We use a simplified sex chromosome model (*35*) that includes a sex determining locus at one end and *n*_*L*_ = 500 coding genes *G*. The expression of each gene is controlled by a *cis*-regulator *C*, and *two* trans-regulators *T*_*m*_ and *T*_*f*_ each of which is expressed either in males or in females. We assume that *G* and *C* sequences are uniformly spaced along the sex chromosome, with adjacent genes *G* recombining at a rate *R*_*g*_ (initially in both sexes), and each *C* regulator being closer to the *G* gene it regulates (at recombination rate *R*_*c*_, *R*_*c*_ < *R*_*g*_, Fig S6). Our simulations use *R*_*g*_ = 0.0005 (resulting in an initial overall map length of 25 cM), and *R*_*c*_ = *R*_*g*_/10. Trans-regulators, such as transcription factors, are unlinked to their target genes, and influence expression on both homologs, whereas c*is*-regulators, such as enhancers, affect expression only of the gene carried on the same chromosome as themselves (*33*). Trans-regulators with sex-limited effects are necessary for dosage compensation to evolve (*35*). For simplicity, we assume that these *T* loci are autosomal.

#### Inversions and reversions

We assume that inversions occur on the Y at a rate *U*_*inv*_. We only consider inversions that include the sex locus (other inversions are not relevant to the topic investigated here as they are not confined to males). We denote *z* the non-recombining fraction of the Y (*z* is therefore comprised between 0 and 1). This variable is also used to measure the endpoint of each inversion on the map. When *z* = 0, X and Y chromosomes recombine freely, but otherwise X-Y recombination only occurs within the chromosomal segment [*z*, 1]. When *z* = 1, the X and Y do not recombine at all. When a new inversion occurs, its size is drawn as a uniform fraction of the non-recombining part of the Y. Specifically, on a Y where recombination is already stopped between 0 and *z*_*i*_, the arrest of recombination will extend, after the new inversion *i*+1 to *z*_*i*+1_ = *z*_*i*_ + (1 − *z*_*i*_)*u*, where *u* is a uniform deviate between 0 and 1. Finally, we assume that reversions can also occur, at a rate *U*_*rev*_, removing the last inversion on the non-recombining part of the Y (we use *U*_*inv*_ = *U*_*rev*_ = 10^−5^).

#### Regulatory traits

The effects of alleles at the *cis*-(*C*) and *trans*-regulators (*T*_*m*_, *T*_*f*_) are modelled as quantitative traits, with Gaussian mutations, denoted by *c, t*_*m*_, *t*_*f*_, respectively. These regulators control allele-specific expression as well as the overall level of expression *Q* of each gene. Mutations in *cis* and *trans* regulators are assumed to occur at rates *U*_*c*_ and *U*_*t*_, respectively, and add a Gaussian deviate to allelic values for these traits (*c* + *dc*∼*N*(0, σ_*c*_), *t* + *dt*∼*N*(0, σ_*t*_)). We use σ_*c*_ = σ_*t*_ = 0.2 and *U*_*t*_ = *U*_*c*_/2 = 10^−4^. Negative trait values are counted as zero. These values are used to compute the total and allele-specific expression values for each coding gene *G*, as explained below. Note that we introduced two trans-regulators per gene, one expressed in males, and the other in females. We could instead assume a single trans-regulator determining two independent traits in males and females, which would be equivalent. The key is that sex specific trans-regulation is possible, so that dosage-compensation can occur. Some genes may be unable to quickly evolve sex-specific regulation. If included, they would not diverge between X and Y, and they would therefore not contribute to recombination arrest.

#### Allele-specific expression and dominance

Arbitrarily denoting with a 1 or 2 subscript two alleles at a gene locus *G*, we assume that the fraction of the protein expressed from allele 1 is ϕ_1,*i*_ = *c*_1,*i*_T(*c*_1,*i*_ + *c*_2,*i*_). This ratio measures the degree of allele-specific expression. With ϕ = 1/2 (i.e. with equally strong cis-regulators on both homologs), alleles are co-expressed, while a departure from 1/2 indicates that one allele is relatively more expressed than the other. The dominance of deleterious mutations occurring on the gene *G* depends on this allele-specific expression and is given by

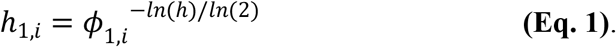

where *h* is a parameter measuring the dominance of the fitness effect of deleterious mutations in a heterozygote when both alleles are equally expressed (with ϕ_1,*i*_ = 1/2, we have *h*_1,*i*_ = *h*). We set *h* = 0.25, corresponding to the average value observed across species (*41*).

#### Deleterious mutations on genes

Deleterious mutations occur on genes *G* at a rate *U*_*G*_ per gene. Their fitness effect *s* is drawn from an exponential distribution with mean *s*_*mean*_. We use *s*_*mean*_ = 0.05. The effects of multiple mutations in the same gene are assumed to be additive, but can be cumulated only up to a maximum effect per gene, *s*_*max*_ (measuring the fitness effect of the gene knock-out). Their dominance depends on the strength of their associated *cis*-regulator. The more they are expressed (relative to the other allele), the larger their effective dominance, as explained above. The fitness effect resulting from the presence of deleterious mutations in gene *i* is

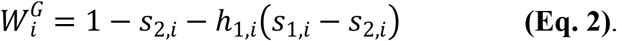

where, by convention subscript 1 denotes the most deleterious allele of the two present in a given individual for that gene *i*.

#### Stabilizing selection on expression levels

We assume that the overall expression level of coding genes is under stabilizing selection with an optimum value *Q*_*opt*_. In males, the total expression level *Qi* equals 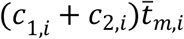, where 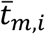 is the average strength of the *trans*-regulators expressed in males, which assumes that both *cis*- and *trans*-regulators are essential for proper expression (neither can be zero). Symmetrically, it is 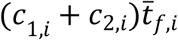 in females. We assume that ln(*Qi*) is under Gaussian stabilizing selection around ln(*Q*_*opt*_) (with *Q*_*opt*_ = 2). We use log-scale to ensure that, irrespective of the intensity of stabilizing selection, the fitness effect of complete regulatory silencing (*Q*_*i*_ = 0) would be *s*_*max*_, the maximum possible fitness effect of deleterious alleles on the coding gene, which we assume to be the same as the effect of a gene knock-out. Denoting by *I* the intensity of stabilizing selection on the expression level, the fitness resulting from the departure from optimal dosage 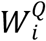 is

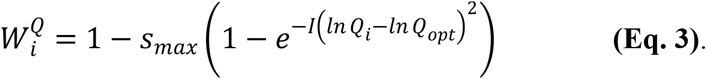

This function is equivalent to assuming that fold-changes in expression levels are under symmetric stabilizing selection, while selection on expression levels *Q*_*i*_ is asymmetric. Unless otherwise stated, we use *I* = 0.1.

#### Individual fitness

Individual fitness is contributed by two components: the fitness consequences of carrying deleterious mutations in the coding gene (whose dominance depends on allele-specific expression), 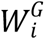, and stabilizing selection on overall expression level 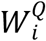. The overall fitness of an individual is computed as the product over all genes *i* of 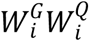.

#### Sexual dimorphism

It is important to note that our model does not include traits selected to be different in males and females. All the genes we consider have exactly the same expression optimum in males and females and we only consider deleterious mutations on genes that have the exact same effect in males and females. The presence of a sex-determining locus and sex-specific trans-regulators implies the existence of some sexually dimorphic traits coded somewhere in the genome, but these traits are absent from the simulations and therefore play no role in the results.

#### Life cycle and simulations

The different events of the life cycle occur in the following order: diploid selection, meiosis with recombination, mutation, and syngamy. Simulations are initialized with *N*_*pop*_ individuals, no polymorphism present, fully recombining sex-chromosomes and optimal gene expression levels (no deleterious allele, all *c* and *t*_*m*_, *t*_*f*_ alleles fixed to 1, all Y chromosome with *z* = 0). After a burn-in phase, mutations producing inversions and reversions are turned on and we follow the dynamics of the system. Unless otherwise stated, we use *N*_*pop*_ = 10^4^.

#### Measures

At regular time steps, we record average regulatory trait values in the population, mean fitness of males and females, average effect of deleterious mutations on the Y (*s*_*Y*_), average dominance of mutations on the Y (*h*_*Y*_), *P*_*silent*_ the probability that ϕ_*Y,i*_ becomes close to zero (below 0.01) so that alleles on the Y become nearly entirely recessive (averaged over all genes), *P*_*dead*_ the probability that deleterious mutations on Y gene copy have reduced fitness by an amount *s*_*max*_, indicating that the gene has entirely degenerated on the Y (averaged over all genes), and the average length of the non-recombining portion of the 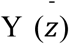. We also record average “sex-switched” fitness, i.e. the average fitness of females computed replacing one of their X by a randomly drawn Y, and the average fitness of males computed replacing their Y by a randomly drawn X (relative to female and male average fitnesses, respectively).

We record all inversions occurring in the population (time of occurrence, start and end point, *s*_*Y*_, *h*_*Y*_, frequency, average regulatory traits, marginal fitnesses). We compute the marginal fitnesses of inversions (denoted *W*_*margY*_) as the product of 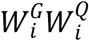 for all genes carried by this inversion (averaged over all Y chromosomes carrying this inversion), relative to the same product computed over all Y chromosomes not carrying this inversion. This quantity indicates whether inversions involve segment of the Y chromosome that present a higher or lower fitness effect compared to the equivalent non-inverted Y segment in the population. We also compute this marginal fitness relative to the average fitness effect of the same chromosomal segment sampled from an X chromosome, and placed in a male carrying the inversion (we use 1000 such samples to compute this value). Indeed, when a reversion occurs, followed by recombination events, it creates new Y chromosomes including (part of) this homologous X segment. We denote this quantity *W*_*margX*_. It measures the average fitness of recombinant Y, relative to the actual Y, and therefore whether reversions could be selectively favored (if *W*_*margX*_ < 1). When *W*_*margX*_ >>1, it indicates that reversions and recombinant Y have a much lower fitness than current Y in the population, which is the signature of ‘stabilized inversions’.

## Supplementary Material

### 1. Some limitations of current models

#### Recombination arrest

Current theory for Y recombination arrest is mostly based on the idea that the evolution of sexually dimorphic traits involves sexually antagonistic loci (SA-loci), where mutations occur that are beneficial in one sex, but deleterious in the other. If SA-loci are widespread, some would inevitably occur on the sex-chromosomes, which would then favor the evolution of tight linkage with the sex-determining locus. There is ample evidence for sexual conflict in general, making this assumption plausible. However, the role of SA-loci in Y recombination arrest is not demonstrated empirically (*3, 5, 26, 27*). This demonstration is inherently difficult to make because SA-loci are difficult to detect and because they can occur after the recombination arrest. Hence, whether this theory explains why sex chromosomes stop recombining is still entirely open.

One limitation of this hypothesis is that sex-linkage is only a particular way to solve intralocus sexual conflicts. Other resolutions based on regulatory evolution can occur as well (*42*– *44*), and the question is really whether these resolutions are more or less likely to evolve than sex-linkage. (a) In the simplest case, the sexual conflict is on the level of expression of a given protein. In this case, sex-specific trans-regulators can easily adjust optimal expression in both sexes, solving the conflict. (b) In a second case, a particular protein modification is favourable in one sex but needs not to be expressed in the other. This locus can evolve to be expressed only in the sex where it is favourable, by changing either cis or sex-specific trans-regulators, solving the conflict. (c) In a third case, two versions of a protein are favourable, each in one sex, but this protein needs to be expressed in both sexes. Here, the conflict can be solved by evolving an heterogeneous gene duplication with each copy becoming expressed in the sex where it is favourable (“subfunctionnalization” scenario). These genetic changes can evolve rapidly. Evolution of sex-specific expression is rapid (*42, 45*) and evidence for SA-polymorphism is limited compared to the signature of resolved expression differences (*43*). Even the most constrained case (c) may evolve rapidly, as demonstrated by the astonishing diversity of newly emerging heterogeneous gene duplication that have been documented at extremely short time scales in some cases (*46*).

Our model also suggests other limitations to the sexual antagonism theory. It is possible to propose a variant of our model where the initial fixation of inversions is promoted by the presence of a SA-locus. However, this appears unnecessarily complex, as such inversions will inevitably occur and fix without those SA loci by capturing “lucky” chromosomal segments with fewer deleterious mutations, which is a very potent mechanism. Then, regulatory sex-antagonistic effects will inevitably emerge if early dosage compensation is allowed, as we show, abolishing the need for SA locus. Finally, even if recombination suppression is caused by the presence of a SA locus, it may not be robust to the occurrence of reversions. The protection of the SA locus against reversion would last only to a point: until the fitness decay caused by degeneration becomes larger than this SA effect. This limitation is not operating with the regulatory model we propose as the protection grows with time (by the accumulation of dosage compensation and its growing “protecting” sex-antagonistic effect).

#### Degeneration

Models about sex-chromosome degeneration are based on selective interference once recombination is stopped. A limitation of these models is that they tend to be inefficient in large populations or on small non-recombining regions, and especially if only deleterious mutations are considered (*14, 15, 47*). Selective interference is inevitable in absence of recombination, but it may not be the only process causing degeneration. We previously showed that cis-regulatory divergence after recombination arrest could efficiently lead to Y silencing and degeneration, in absence of selective interference (*35*). This process works faster and can explain degeneration of very small non-recombining regions. Here too, there is ample evidence that selective interference occurs, but there is currently no evidence that it is the only mechanism at work, especially in species with large population sizes and small degenerated strata.

#### Dosage compensation

Dosage compensation (DC) tends to be considered separately from the process of degeneration and always assumed to evolve late, as reported in virtually all papers and textbooks since Ohno’s 1967 book (*23*). This claim is based on the causal chain of events: in current theory, DC results from degeneration, not the other way around. In fact, degeneration and DC are likely to be nearly simultaneous, at least for dosage sensitive genes, in any viable theory, as otherwise males would suffer an unbearable fitness decline relative to females during Y degeneration. However, models incorporating degeneration and DC tend to be currently missing, with some exceptions (*14, 39*). We recently showed that silencing and DC could theoretically occur slightly before and cause rather than result from degeneration (*35*).

### 2. Inversion and reversion assumptions

#### Inversions

The model assumes Y-carried mutations that stop recombination on a portion of the Y (and that this non-recombining region can be later extended or reduced). It helps to think that these mutations correspond to inversions, but there is really nothing ‘inversion-specific’ in the model. Chromosome collinearity is not altered in the code, and there is no specific assumptions determining how recombination proceeds on non-perfectly collinear chromosomes. The non-recombining region is determined using the variable *z*. Recombination does not occur between 0 and *z* and occurs between *z* and 1. “Inversion” mutations only alter this *z* value, but do not specify the exact mechanism by which this recombination arrest actually takes place. Hence, the results presented extend to other mechanisms of recombination arrest, as long as they involve a genetic change occurring on the Y (e.g. heterochromatinisation, hotspot presence etc.). In addition, the mutations considered here may not represent “real” inversions. Large real inversions do not necessarily create complete linkage across the inverted region, as double crossovers may be possible. Thus, the inversion that we consider would correspond to the non-recombining part of real inversions. This issue is however minimized by the fact that large inversions do not contribute to recombination arrest in our model (Fig S1C, S2C).

#### Reversion

We introduced reversions to make sure recombination arrest is not unescapable. Without this, inversions would inevitably occur and fix quickly, and no specific explanation would be required for Y recombination arrest. Such a model would be rather superficial, since the outcome would directly results from this hypothetical constraint. A similar mistake was made in early sex-chromosome models, which assumed that deleterious mutations could only occur on the Y and then concluded that the Y would inevitably degenerate.

We considered another model for reversion, where each reversion was restoring recombination on all the Y, not just canceling the last inversion. This other model lead to similar results suggesting that it does not matter much how precisely recombination can be restored, as long as it can. This other model involved however more complex dynamics. One of its specificity is to make reversions less and less likely as the Y degenerates, by introducing a strong dependence between successive strata. In particular, a new inversion can be protected from reversions by the sex-antagonistic effects accumulated on other previous strata. This model also introduces coupled reversion-inversion dynamics, since a reversion can only persist if it is immediately followed by an inversion that stops recombination on the part of the Y that is already degenerated.

Other model of reversions could be imagined, but most other solutions would be computationally challenging to perform. For instance, reversion breakpoints could be randomly sampled anywhere in the non-recombining region. Such an approach would create each time a new inversion (with a new starting and ending point). Since we need to keep track of all inversions, this adds an important computational burden, without adding any interesting process. Another, very complicated way to allow for recombination restoration, would involve inversion on the X. If an inversion is present on the Y, but an inversion also occurs on the X, it could partially restore recombination, but this model would be extremely complex to run and analyze as it would require to keep track of X and Y collinearity and introduce complex additional assumptions to decide how recombination proceeds on non-perfectly collinear chromosomes. It would also require doing very complex bookkeeping of all possible chromosome orderings, including inversions that do not include the sex-determining locus.

### 3. Comparison to current theory and empirical observations

Comparisons of different theories can be made on several grounds that we can broadly categorize in terms of plausibility, parsimony and predictive power.

#### Plausibility

The model we propose involves a very general process with almost no specific assumption. All ingredients are basic genetic features of eukaryotes, such as deleterious mutations, *cis* and *trans*-regulators, inversions, stabilizing selection on dosage, and partial dominance. All these features have been extensively demonstrated. It also includes more specific assumptions, such as the occurrence of genetic mutations that can alter recombination rate on the Y, in a reversible manner. There is ample evidence for genetic variation in recombination rates, and this is an ingredient that must be present in any theory aiming at explaining Y recombination arrest. As discussed above, our way to represent reversion is certainly a drastic simplification, yet other models of Y recombination arrest do not even include the possibility of a restauration of recombination, especially once degeneration has started to occur. As we show in the model without regulatory evolution, if this possibility is included, degeneration would not occur on the long run, as reversions allow eliminating partially degenerated Y from the population. The presence of a sex-antagonistic polymorphic locus is also unlikely to be sufficient to counteract this process. The model we propose naturally generates ever increasing sex-antagonistic effects, which accumulate over all dosage sensitive genes, and naturally increase with the degree of degeneration. Hence, this mechanism is more likely to consistently favor the maintenance of recombination suppression than the occurrence of a handful of isolated loci with sex-antagonistic effects. Models allowing for dynamical accumulation of sex-antagonistic alleles might be possible, but they have not been worked out, and probably demand that an unrealistic large fraction of genes is involved in sexual dimorphism. Our model distinguish a proximal and ultimate cause for recombination arrest. The origin and maintenance of recombination suppression have distinct causes. Initially, there is no selection against recombination. However, a fortuitous but selective phenomenon (lucky inversion spread and fixation) causes a selection pressure against recombination that was previously absent (the quick emergence of regulatory sex-antagonistic effects), and which protect this inversion from being subsequently lost. Distinguishing the problem of origin and the maintenance of a trait has often been considered in other contexts, notably the evolution of sex (48).

#### Parsimony

The theory we propose tends to be more parsimonious than current theory. It makes loci with sex-antagonistic effects superfluous, while all the ingredients present in our model are required in any global theory of sex chromosomes. For instance, deleterious mutations are necessary for degeneration, sex-specific regulatory changes are necessary for dosage compensation, and mutation altering recombination are necessary to explain recombination suppression. It is also parsimonious as it explains all the process of Y recombination arrest, degeneration and dosage compensation in a single model where all steps are integrated and work consistently within the same set of assumptions. In contrast, current theory is mostly made of series of models addressing each step separately, with different sets of assumptions.

#### Predictive power

Compared to current theory, our theory explains the same global pattern seen across many eukaryotes. Y or W chromosomes are often non-recombining, degenerated and at least partially dosage-compensated. Even if the causal explanation for each of these steps differs between current theory and the theory presented in this paper, observations distinguishing them may not be available yet. The “established model” lacks decisive empirical support, despite decades of investigations. In particular, there is no firm evidence that sex-antagonistic loci cause recombination suppression. However, absence of evidence is not evidence of absence, and the mechanism works in principle. There is ample evidence for the existence of male female antagonistic selection, indicating that this explanation could work. In comparison, there is no indication that early dosage compensation on dosage-sensitive genes generates sex-antagonistic effects on young sex chromosome, but this has not been looked for. There are indications of early evolution of dosage compensation or regulatory evolution in some species. This is an essential piece of information, consistent with our theory, but not proving that recombination arrest is caused by these modifications. Current theory does not predict a particular size for Y chromosome strata. It is however difficult to explain that small strata degenerate by selective interference if they contain only few genes. Our theory tend to indicate that rather small strata are involved in recombination arrest, although the exact size depends on details that we did not investigate. For instance, if there is an important heterogeneity for dosage sensitivity among genes, strata size could be partly dictated by the chance localization of genes that are strongly dosage sensitive. In any case, our theory can certainly explain better why small strata can occur and degenerate. It may also easily explain cases where divergence appears nearly continuous along the X-Y chromosome pair, as if many small strata accumulated. Conversely, if several small strata occur on a short time interval, it may look as if a single large stratum evolved (see Fig S3 for the heterogeneity of simulation replicates). Several but not all strata result from inversions. Our model is presented using the term “inversion”, but as explained in sup. mat. 2 there is nothing inversion-specific in our model, and other genetic modification suppressing recombination would work.

Empirically, the relative timing of degeneration and dosage compensation is not easily established. Dosage insensitive genes will almost never evolve dosage compensation, by definition, unless they are caught in a chromosomal level mechanism. Hence seeing that some genes are degenerated but not dosage compensated is not very informative. It may simply indicate that they are dosage insensitive, not that dosage compensation evolves after degeneration for dosage sensitive genes. Hence, observing that degeneration is more advanced than dosage compensation is not refuting our theory. Similarly, our model does not require complete early and full dosage compensation of all genes in a stratum. Only dosage sensitive genes are expected to evolve quick and early compensation, so observing partial or gene specific compensation is not refuting what we propose. The evolution of dosage compensation in neo-Y systems, after the fusion of an autosome to an already existing Y might be special cases, where a chromosomal-level dosage compensation mechanisms is just extended, without having to evolve from scratch. Such chromosomal-level mechanism would involve all genes (dosage sensitive or not) and therefore lead to a quite specific patterns. Other observations consistent with our theory include the absence of degenerated sex chromosomes in species where sex chromosomes are expressed in haploids, as in *Ectocarpus* algae (*49*). The presence of evolutionary strata on young mating-type chromosomes in species entirely lacking sexual dimorphism (*50*). could be explained by regulatory evolution if mating-type specific expression could evolve. However, recombination may evolve for other reasons in species with specific reproductive modes (*51*).

Our model predicts that Y recombination arrest and degeneration should be quicker in large populations, everything else being constant. This pattern may not hold, or may be saturating for larger population sizes than the ones we considered. This is open for investigation, but would require very long computation time. Empirically, we do not have clear indication of the effect of population sizes on the patterns of sex-chromosome evolution. There are many confounding factors (shared ancestral sex determination systems across species, different ages of sex chromosomes, the initial recombination rate around the sex-determining locus, and the particular case of achiasmate species), but this is an interesting avenue for future research.

**Fig S1.**
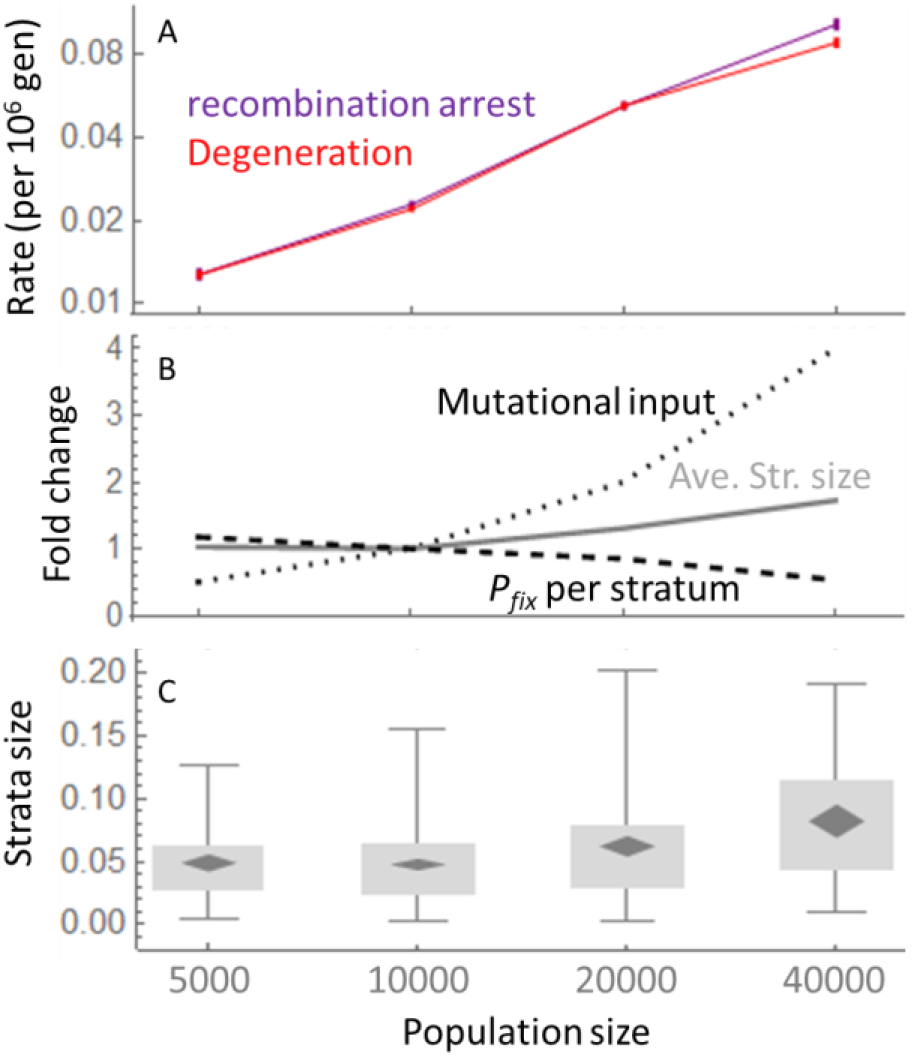
Effect of population size on Y recombination arrest and degeneration. **(A)** Rate of recombination arrest (fraction of Y becoming non-recombining, purple) or degeneration (proportion of gene knocked-out, red) per million generations, for different population sizes (*x*-axes). These processes are approximately linear in time (not shown). **(B)** Contribution of the different factors to the variation in the rate of recombination arrest. The absolute number of inversions arising is proportional to population size (dotted curve, scaled relative to the value at *N* = 10,000). Stabilized inversions tend to be larger in larger populations (gray line, scaled relative to the value at *N* = 10,000). The probability that an inversion fixes and becomes stabilized decreases with population size (dashed line, scaled relative to the value at *N* = 10,000). The contribution of these three factors explain the difference in rates shown on panel A. **(C)** Distribution of stabilized inversions sizes for different population sizes. Black diamonds show means and confidence intervals; gray boxes show limits of 25% and 75% quantiles; whiskers show 5% and 95% quantiles.

**Fig S2.**
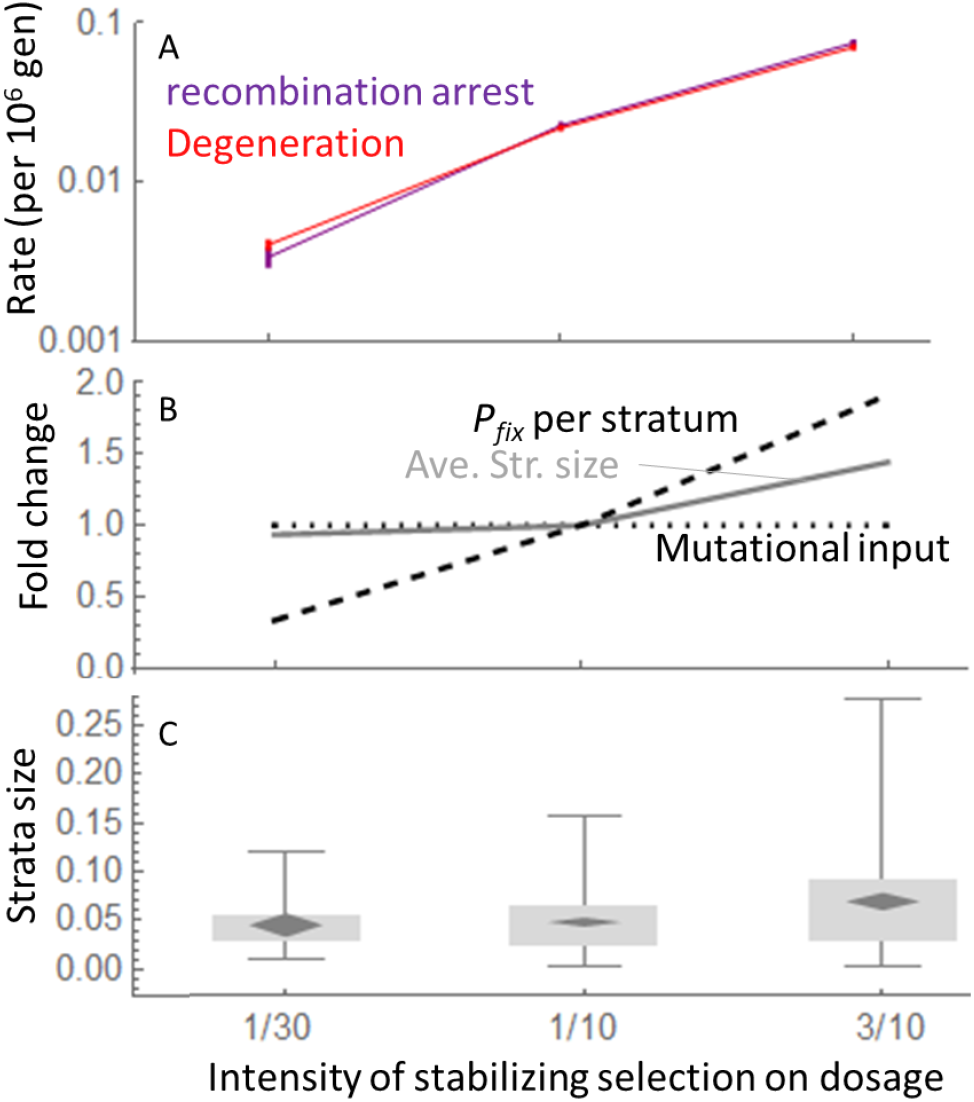
Effect of the intensity of stabilizing selection on gene expression levels, on Y recombination arrest and degeneration. **(A)** Rate of recombination arrest (fraction of Y becoming non-recombining, purple) or degeneration (proportion of gene knocked-out, red) per million generations, for different intensities of stabilizing selection (*I*) on dosage (*x*-axes). These processes are approximately linear in time (not shown). **(B)** Contribution of the different factors to the variation in the rate of recombination arrest. The absolute number of inversions arising is identical for the different values of *I* and correspond to the value at *N* = 10,000 (dotted line). The probability that an inversion fixes and becomes stabilized increases with *I* (dashed line, scaled to the value at *I* = 0.1). Stronger stabilizing selection increases the sex-antagonistic fitness effect of nascent dosage compensation, which increase the chance that an inversion is stabilized and that large inversions escape reversion. The contribution of these three factors explain the difference in rates shown on panel A. **(C)** Distribution of stabilized inversions sizes for different values of *I*. Black diamonds show means and confidence intervals; gray boxes show limits of 25% and 75% quantiles; whiskers show 5% and 95% quantiles.

**Fig S3.**
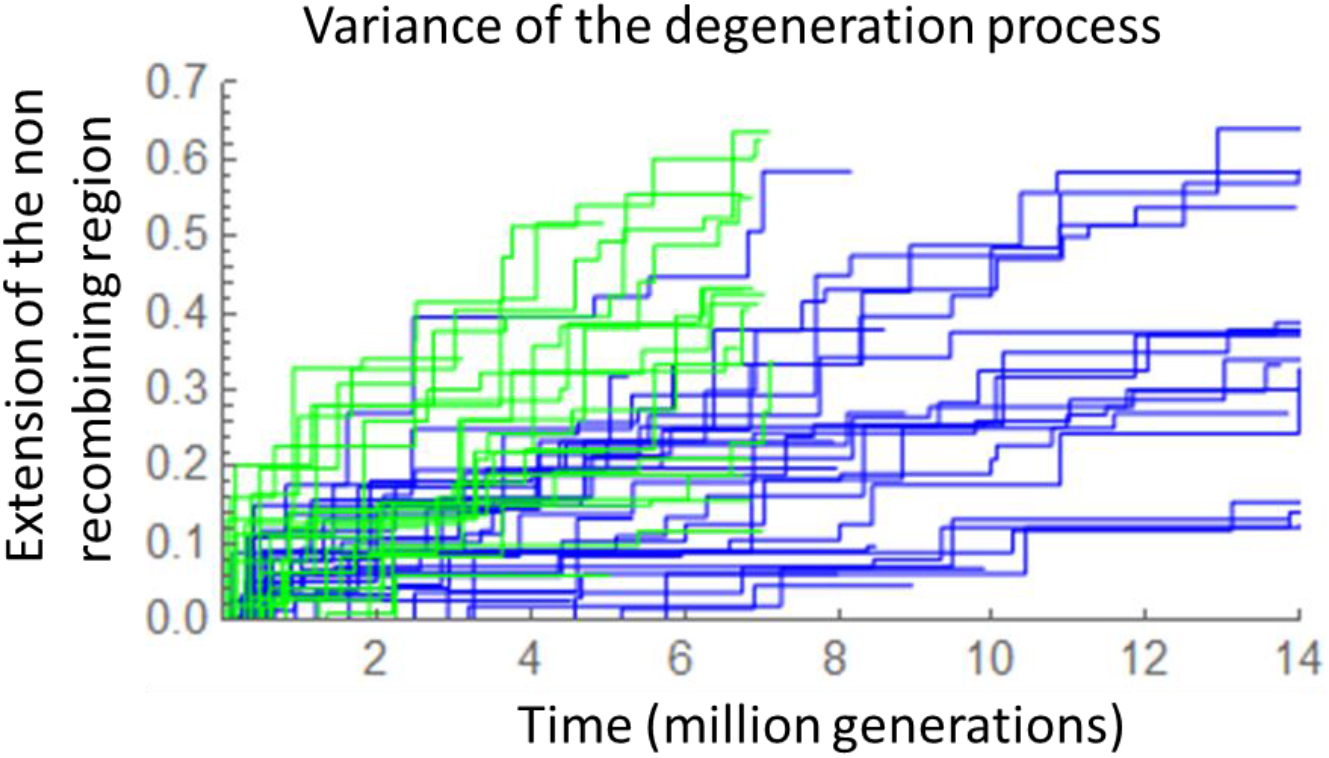
Stairplot representing the occurrence of stabilized strata on Y in different evolutionary replicates. Each line corresponds to the stairplot illustrated in black in Fig. 2A for one replicate. Parameters as in Fig. 2, except for population size: blue *N*_*pop*_=10,000, green *N*_*pop*_=20,000. Note that runs with *N*_*pop*_ = 10,000 and 20,000 were stopped at 7 million generations and 14 million, respectively. In both cases, some runs were slower and were interrupted before reaching this limit, because of computation time limits. The figure shows that the process of recombination arrest is highly stochastic.

**Fig S4.**
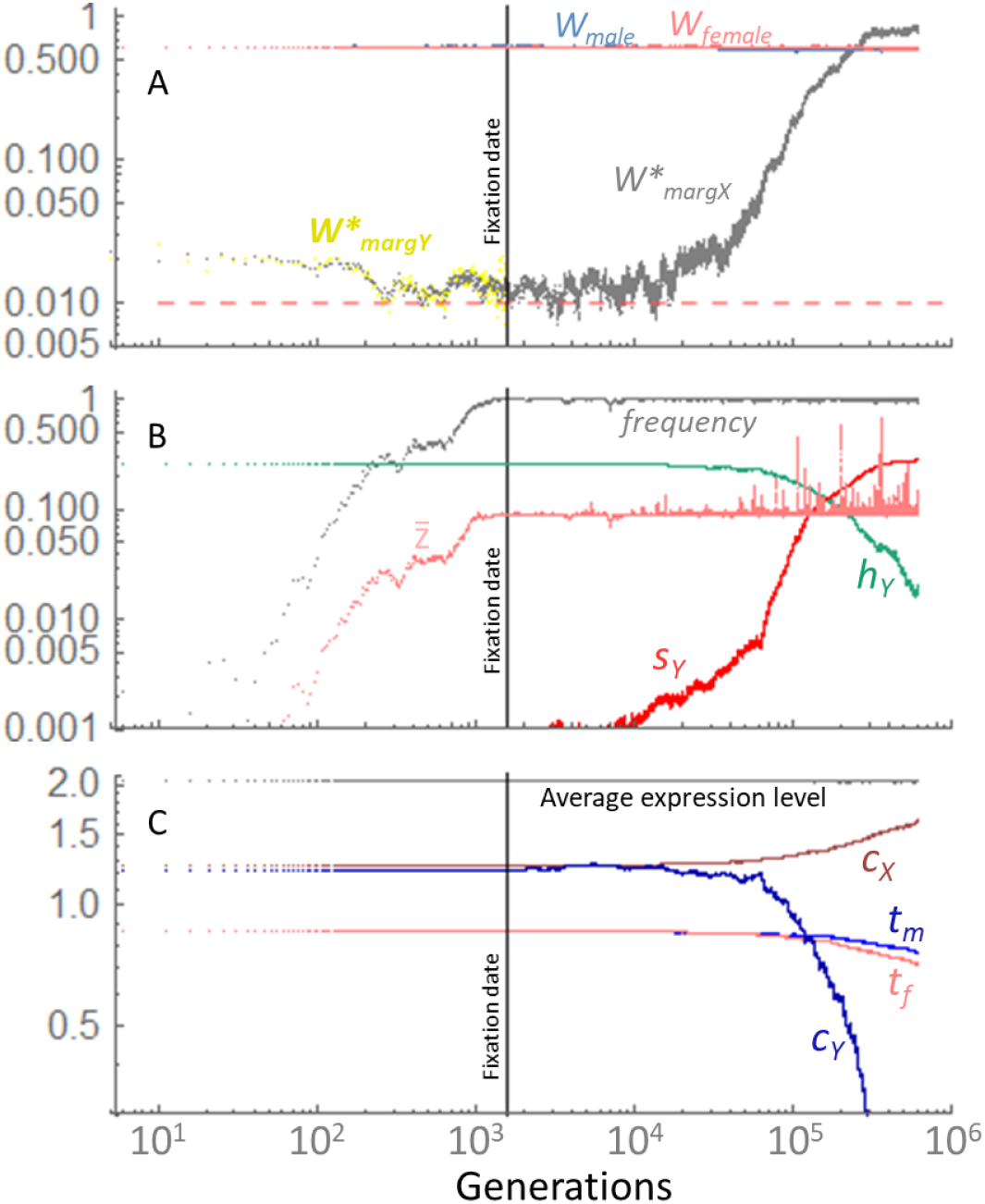
Details of the fixation and stabilization of a first stratum. (**A**) *x*-axis, inversion age: the number of generations since the appearance of the inversion (in log-scale). Gray : marginal fitness of the inversion relative to the same chromosomal segment on the X if it was in a male (*W*_*margX*_, see methods). After fixation, *W*_*margX*_ measures the sex-antagonistic effect of nascent dosage compensation. Yellow: marginal fitness of the inversion relative to the same chromosomal segment among Y-chromosomes not carrying the inversion (*W*_*margY*_, see methods). Note that *W*_*margY*_ cannot be computed after the inversion fixes as all Y-chromosomes carry the inversion. Both these fitness values are represented minus 0.99 (and noted with a *) to allow for a better visualization on the *y*-axis log-scale. Consequently, the red dashed line at 0.01 represents a marginal relative fitness of 1. Average fitness of males and females in the population are also indicated in blue and pink. (**B**) Gray: Frequency of the inversion. Pink average fraction of the non-recombining Y in the population 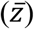. Green: average dominance of deleterious mutations on the inversion (*h*_*Y*_). Red: average deleterious effect of mutations among genes on the inversion (*s*_*Y*_). (**C**) Regulatory trait variation. Dark blue: average *cis*-regulatory trait on the inversion. Brown: average *cis*-regulatory trait on the corresponding X segment. Blue: average *trans*-regulatory trait associated to genes on the inversion. Pink: average *trans*-regulatory trait associated to genes on the corresponding X segment. Gray: average total gene expression per genes (undistinguishable in males or females) for genes present on the inversion. Note that between 10^3^ and ∼5.10^4^ generations, there is enough sex-antagonistic effect of nascent dosage compensation to protect the inversion from reversion, as seen on panel A with *W*_*margX*_, while there is still almost undetectable X-Y *cis*-divergence and male –female trans-regulatory divergence. On all panels, the vertical bar indicates the date at which the inversion fixes for the first time (because of the occurrence of reversions the frequency slightly departs from one afterwards).

**Fig S5.**
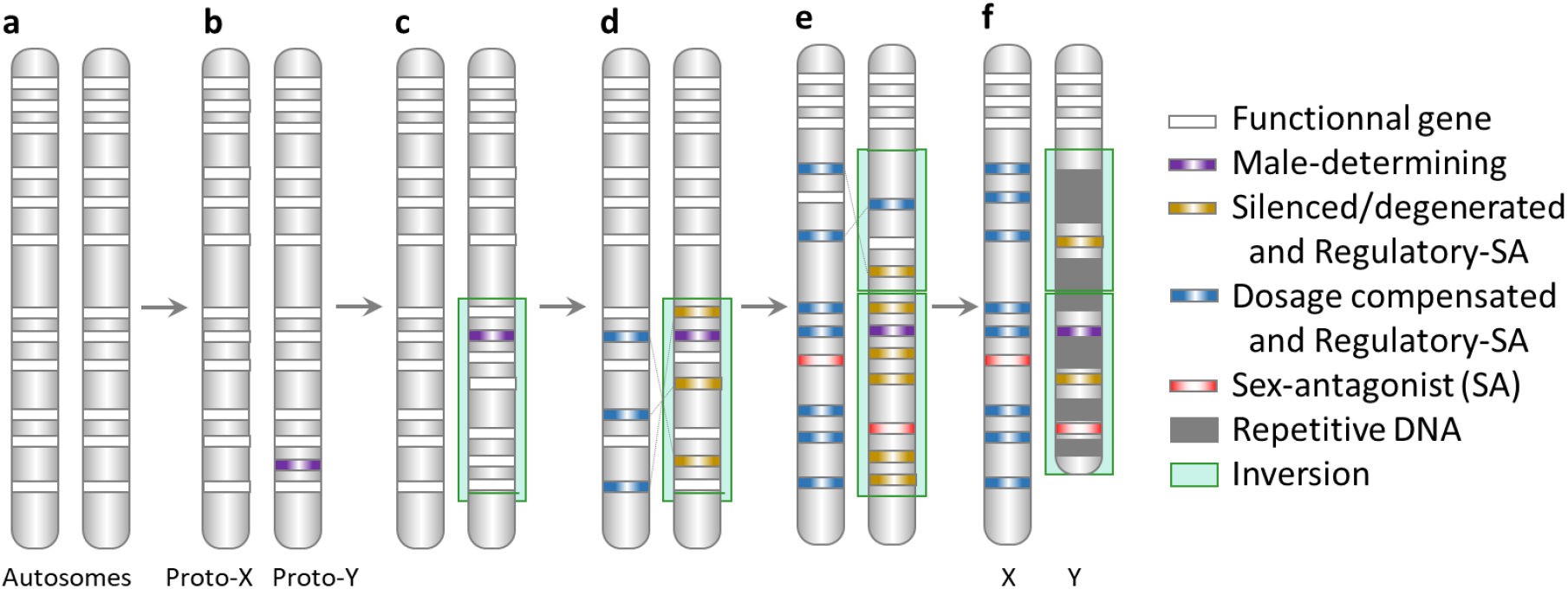
Overview of Y evolution proposed here, inspired from Fig. 1 in (*25*) representing current theory. (**a**) and (**b**) A pair of autosomes acquire a sex-determining locus (purple). (**c**) A lucky inversion carrying fewer or milder deleterious mutations than average selectively fixes in the population (green). (**d**) Cis- and trans-regulatory divergence causes regulatory sex-antagonisitic effects on dosage sensitive genes, which stabilizes the inversion on the long term. Y genes tend to be silenced (yellow) and accumulate deleterious mutation (degeneration by regulatory evolution), while their X copy already show dosage compensation (blue). (**e**) The process repeats itself with another sex-linked inversion, creating another stratum. Some sex-antagonisitic alleles can occur and be maintained on these sex-linked regions, but they were not involved in recombination arrest (red). (**f**) The absence of recombination leads to the accumulation of repetitive DNA and/or structural changes (deletions).

**Fig S6.**
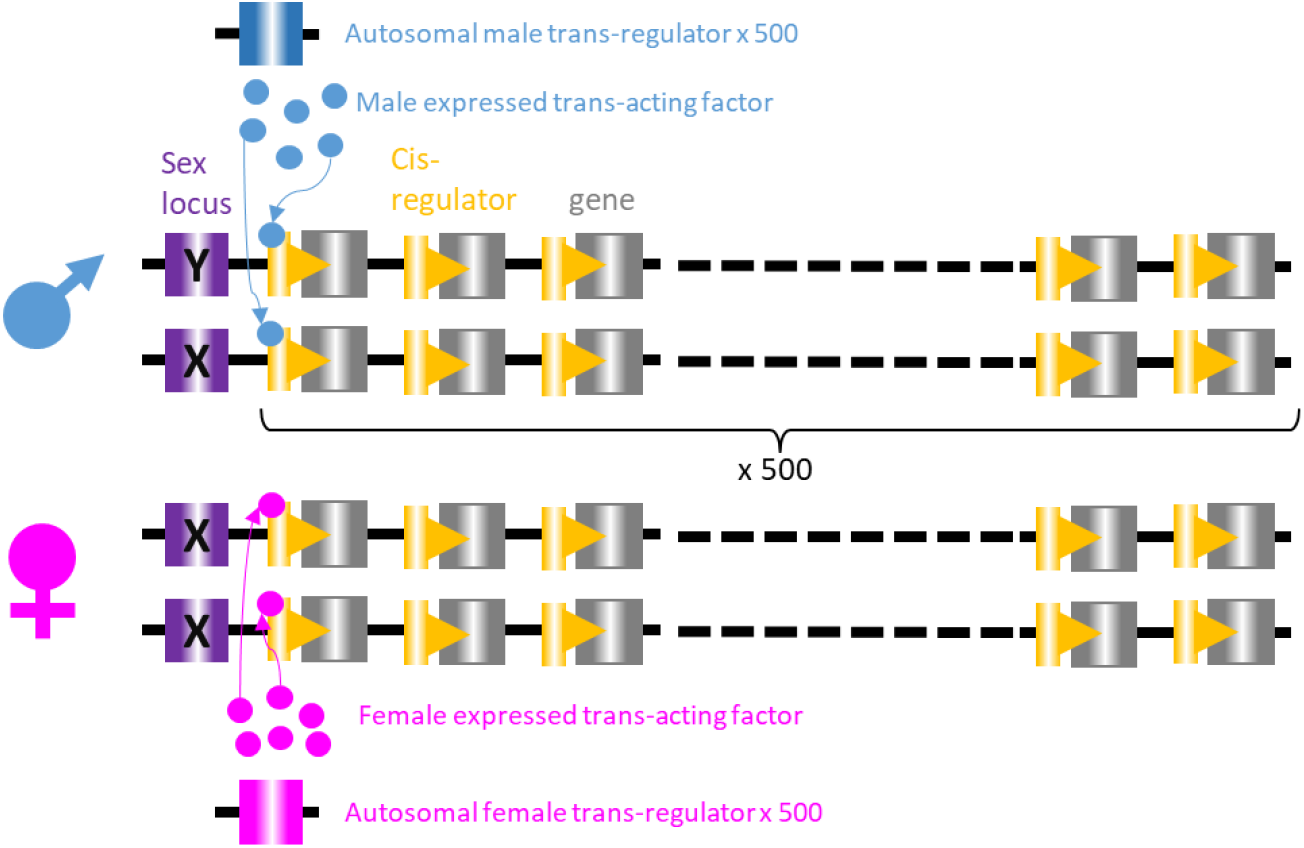
Overview of the genome simulated. The sex chromosome pair carries the sex locus at one end with two alleles (X/Y) determining sexes (XX female, XY male). This chromosome carries 500 coding genes, each with a *cis*-regulator. Each *cis*-regulator interact with a *trans*-acting factor expressed from an autosomal *trans*-regulator which differs in males and females. Genes, *cis*- and *trans*-regulators all mutate (partially deleterious mutations on genes, Gaussian deviations on cis and *trans* regulatory traits). Gene expression level is under stabilizing selection, and dominance of deleterious alleles in coding sequences depend on their relative allele-specific expression. See Fig 1 in (*35*) for visual sketch of expression patterns and selection on each gene.

## Notes

### Competing Interest Statement

The authors have declared no competing interest.

